# A Computational Framework for Extracting Mechanistic Hypotheses from Quantitative Data of Morphological Dynamics

**DOI:** 10.64898/2026.06.10.731458

**Authors:** Koji Kyoda, Hatsumi Okada, Shuichi Onami

## Abstract

Advances in live-cell imaging and image analysis have made it possible to quantitatively measure the spatiotemporal morphological dynamics of biological phenomena at scale. However, a general framework is still lacking for systematically extracting, from the resulting multivariate data, which relationships between phenotypic characters are mechanistically interpretable and which gene perturbations disrupt those relationships. Here, we propose a computational framework for extracting mechanistic hypotheses and candidate genes from quantitative data of morphological dynamics. First, we detect reproducible correlations between phenotypic characters in wild-type data and interpret them as mechanistic hypotheses in light of existing knowledge. Next, we perform outlier analysis on data obtained under gene perturbation and extract, as candidates, genes that selectively disrupt relationships between phenotypic characters maintained in the wild type. We further integrate the extracted relationships into a spatiotemporal network to provide an overview of how phenotypic characters are linked across the developmental process. As a proof of concept, we applied the framework to quantitative data on nuclear division dynamics during early embryogenesis in *Caenorhabditis elegans* and recovered relationships between phenotypic characters consistent with known mechanisms while prioritizing candidate genes. This framework provides a useful basis for efficiently generating testable mechanistic hypotheses from quantitative data of morphological dynamics.

**Author Summary:** How does a single cell give rise to a complex organism? Answering this question requires understanding not only which genes are active, but how the physical behavior of cells—their shapes, positions, and movements—is coordinated across time and space. Live-cell imaging now allows researchers to measure these morphological dynamics in quantitative detail, yet extracting biological meaning from the resulting large, high-dimensional datasets remains a challenge. Here we present a computational framework that addresses this challenge by treating correlations between quantitative morphological measurements as windows into the underlying biological machinery. Applied to the nematode *Caenorhabditis elegans*, a powerful model organism whose early development is exquisitely reproducible, our approach automatically identifies pairs of cellular measurements that reliably co-vary in normal embryos and interprets these relationships as reflecting shared biological mechanisms. When genes are inactivated one at a time and the resulting embryos deviate from the expected co-variation, those genes are flagged as candidates for the disrupted mechanism. In a systematic test using embryos in which 263 genes had been individually inactivated, the framework correctly prioritized genes with known roles in spindle positioning and cell polarity. By converting large-scale morphodynamic datasets into a network of testable mechanistic hypotheses, this framework offers a broadly applicable strategy for moving from quantitative phenotyping to mechanistic understanding across diverse biological systems.

## Introduction

Advances in live-cell imaging, four-dimensional microscopy, and image analysis now make it possible to continuously observe biological processes such as development, tissue morphogenesis, and cellular responses in space and time and to quantify their morphological dynamics. Recent studies have accelerated the accumulation of quantitative data of morphological dynamics, including analyses that model single-cell shape changes over time from live-cell images [1], analyses that extract heterogeneity and common principles of organoid morphogenesis from long-term three-dimensional live imaging [2], and analyses that systematically quantify morphological dynamics at the whole-embryo scale [3]. In addition, image-based profiling / morphological profiling, exemplified by CellProfiler and Cell Painting, is widely used as a framework for extracting large numbers of morphological characters from images and comparing differences induced by perturbations or cell states [4,5,6].

Early *Caenorhabditis elegans* embryos provide an excellent system for testing methodologies for quantitative analysis of morphological dynamics because their cell lineage is highly reproducible [7] and systematic perturbation by RNAi is feasible [8]. Foundational tools for automated cell-lineage tracing and quantitative measurement of nuclear division dynamics have already been established [9,10], and large-scale datasets of RNAi embryos have also been accumulated [11,12,13,14]. These data constitute a valuable resource for computational exploration of the biological mechanisms underlying phenotypes.

Correlation analysis is one of the most fundamental approaches for identifying dependencies among observed variables. In morphology and spatiotemporal imaging, it has been used to evaluate the independence of high-dimensional morphological characters in yeast [15], to propose a control model for cell-cycle timing in *C. elegans* embryos [16], and to analyze functional network architecture across multimodal cortical areas using correlation structures derived from cell-resolved neuronal activity time series acquired by contiguous-wide two-photon imaging [17]. However, a generalized method has not yet been sufficiently established for extracting reproducible relationships between phenotypic characters at scale from quantitative data of morphological dynamics and coupling them with perturbation data to search for mechanistic hypotheses and candidate genes. We therefore propose a computational framework based on correlation analysis and outlier analysis to extract mechanistic hypotheses from quantitative data of morphological dynamics, and we demonstrate its utility using nuclear division dynamics data from early *C. elegans* embryos as a proof of concept.

## Results

### Basic concept of the framework for extracting mechanistic hypotheses

The proposed framework starts from relationships between phenotypic characters that are stably maintained in the wild-type group and extracts the mechanistic hypotheses underlying them. First, phenotypic characters are extracted from data of morphological dynamics in the wild type, and correlations between character pairs are detected. When a reproducible correlation is observed for a pair of characters, it suggests the existence of some real generative process underlying that pattern, potentially involving direct causation, a common upstream factor, geometric constraints, or unobserved factors (Figure 1A). In this framework, such relationships are interpreted in light of existing knowledge to derive mechanistic hypotheses. Furthermore, by detecting samples that deviate from those relationships under perturbation conditions, genes involved in the processes supporting the relevant relationships are extracted as candidates (Figure 1B).

**Figure 1.**
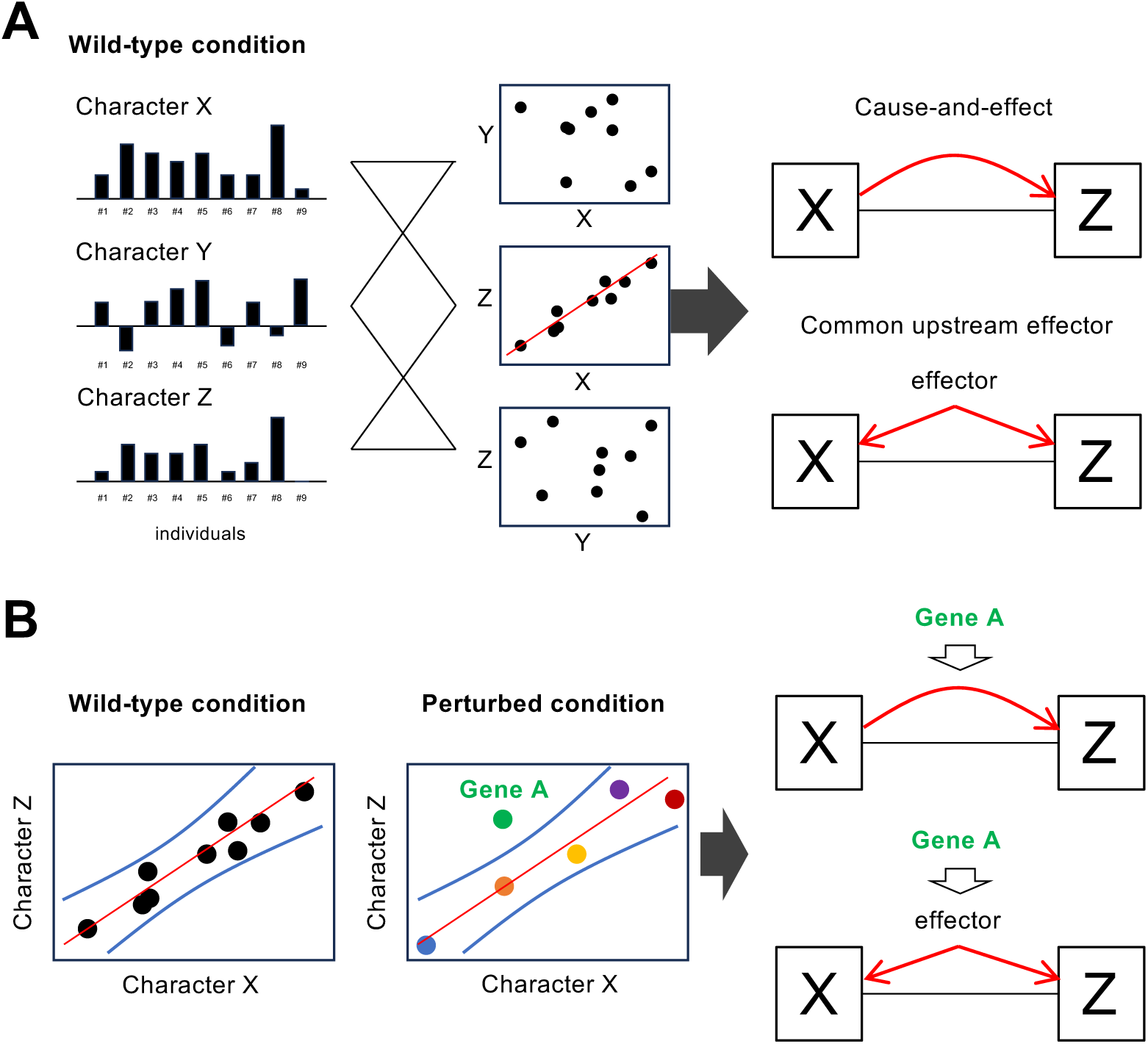
Overview of the framework for extracting mechanistic hypotheses and candidate genes from quantitative data of morphological dynamics. (A) Reproducible correlations between phenotypic characters are extracted from wild-type data, and the shared mechanism underlying them is formulated as a hypothesis. High correlations are likely to reflect some real generative process, including direct causation, common upstream factors, or geometric constraints. (B) When a sample under perturbation conditions falls outside the prediction interval constructed from wild-type data, the perturbation is regarded as having disrupted a relationship between phenotypic characters maintained in the wild type, and the corresponding gene is treated as a mechanistic candidate gene supporting that relationship.

### Detection of correlations in real data

To examine whether stable correlations can be detected between phenotypic characters extracted from data of morphological dynamics, we applied correlation analysis to quantitative data on nuclear division dynamics during early embryogenesis in wild-type *C. elegans* embryos. We first defined intracellular and intercellular characters, including nuclear positions, division orientations, and inter-nuclear distances, and thereby constructed a total of 421 phenotypic characters (Table S1). By grouping characters measured at the same timing, these 421 characters were classified into 46 developmental stages across embryogenesis. We automatically extracted these 421 characters from 59 wild-type embryo datasets and obtained a phenotypic expression profile for each embryo.

Next, from the phenotypic expression profiles of the 421 characters in wild-type embryos, we calculated the correlation coefficient r for every pair of characters. Extracting pairs satisfying |r| > 0.5 yielded 1,824 correlations among 388 phenotypic characters (Table S2). This threshold enables extraction of moderate or stronger correlations between characters [18]. Among them, 4% (73/1,824) were very strong, 14% (249/1,824) were strong, and 82% (1,502/1,824) were moderate correlations (Figure 2A). These results show that large numbers of reproducible relationships between phenotypic characters can be extracted from quantitative data of morphological dynamics in early embryogenesis.

**Figure 2.**
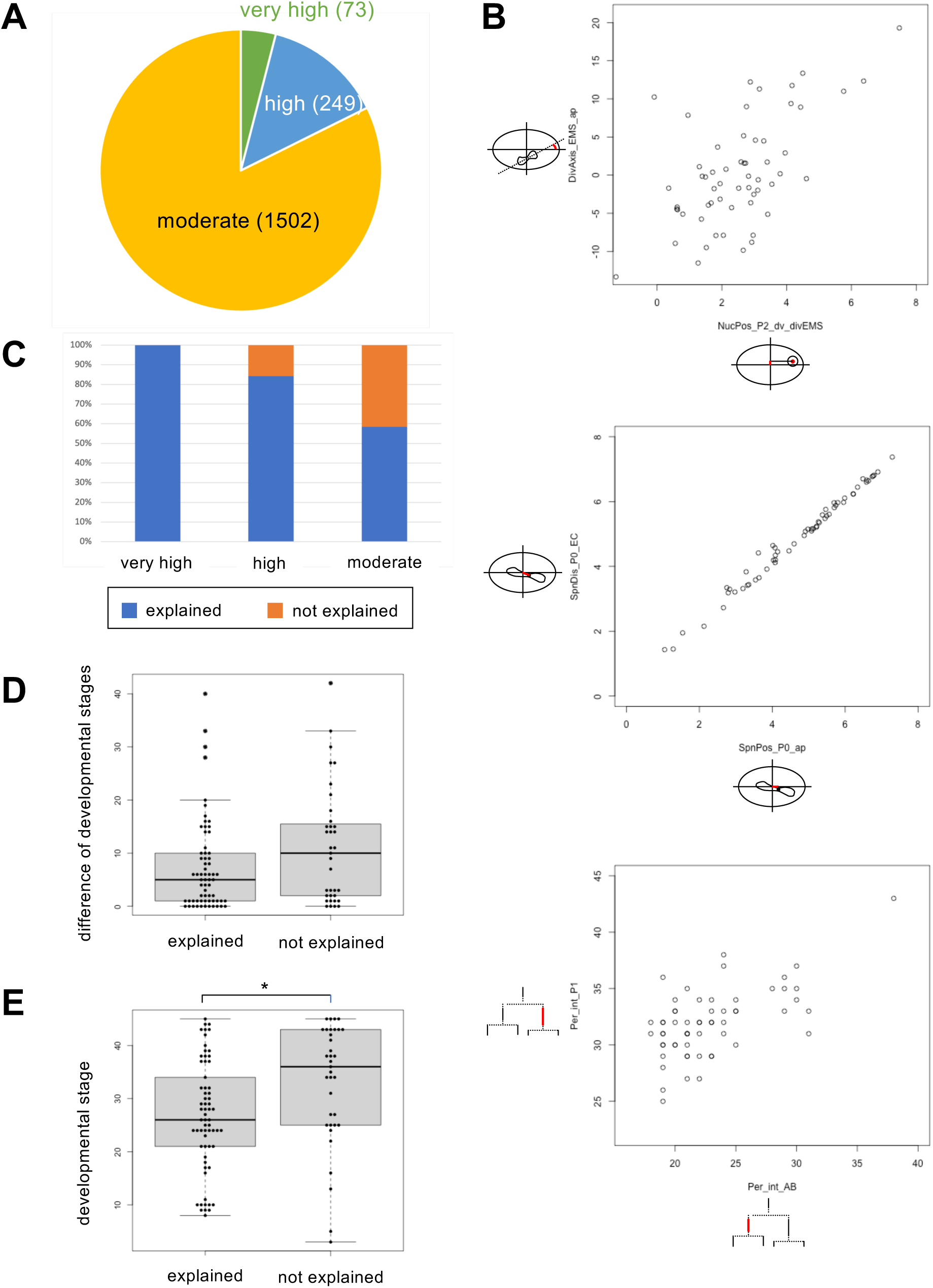
Results of mechanistic hypothesis extraction by correlation analysis. (A) Breakdown of the 1,824 detected correlations by correlation strength. Values in parentheses indicate the correlation coefficient ranges. (B) Three representative examples consistent with known mechanisms. Upper panel: spindle position of the P_0_ cell along the anterior-posterior axis and distance from the embryo center. Middle panel: dorsoventral position of the P_2_ nucleus and orientation of the EMS division axis. Bottom panel: interphase length of the AB cell and that of the P_1_ cell. (C) Proportion of relationships explainable by existing knowledge, stratified by correlation strength. (D) Distribution of developmental-stage differences for character pairs in the explainable and unexplainable groups. (E) Distribution of the later developmental stage within character pairs in the explainable and unexplainable groups.

To evaluate the possibility that the detected relationships were merely incidental high correlations, we generated pseudo phenotypic expression profiles and performed the same analysis. Specifically, for each character, the 59 values were independently randomly shuffled so that the range of each character was preserved while only the actual correspondence between characters was broken. In this pseudo profile, no correlation coefficient exceeding the threshold (|*r*| > 0.5) was observed except for six pairs (Table S3). Therefore, almost all high correlations detected in wild-type embryos are unlikely to be numerical fluctuations arising by chance and instead appear to reflect substantive relationships between phenotypic characters.

### Validation of the framework for extracting mechanistic hypotheses

To show that the framework can recover relationships consistent with known biological mechanisms, we highlight three representative examples among the detected correlations. The first example is the relationship between the spindle position of the P_0_ cell along the anterior-posterior axis and the distance from that spindle position to the embryo center (upper panel of Figure 2B). As the P_0_ spindle approaches the embryo center, both its coordinate along the anterior-posterior axis and its distance from the embryo center change. Although this relationship may appear geometrically trivial at first glance, it actually arises from a polarity-dependent process in which the P_0_ spindle is formed along the anterior-posterior axis and subsequently displaced toward the posterior pole. Thus, this example shows that the correlation between the two characters is supported by the establishment and maintenance of cell polarity and by force-generating mechanisms acting on astral microtubules [19].

The second example is the relationship between the dorsoventral position of the P_2_ nucleus and the angle between the EMS division axis, projected onto the anterior-posterior-dorsoventral plane of the embryo, and the anterior-posterior axis of the embryo (middle panel of Figure 2B). As the P_2_ nucleus is positioned more dorsally, the angle between the spindle-formation axis of the EMS cell and the embryo anterior-posterior axis increases. In early *C. elegans* embryos, EMS spindle orientation is known to depend on a signal from the P_2_ cell, such that the EMS spindle points toward the P_2_ side [20]. Because the P_2_ nucleus is located near the cell center, nuclear position can be regarded as a representative indicator of P_2_ cell position. This relationship can therefore be interpreted as reflecting the signaling mechanism from P_2_ to EMS.

The third example is the relationship between the interphase lengths of the AB and P_1_ cells (bottom panel of Figure 2B). A clear correlation was observed between the interphase lengths of these two cells. Rather than implying that AB directly controls P_1_, this likely reflects shared factors such as temperature or the overall developmental rate of the embryo. Indeed, the timing of early cell divisions in nematode embryos has been reported to be temperature dependent [21,22]. This example shows that the framework can capture not only direct causal links but also mechanistically meaningful relationships supported by common upstream conditions.

### Evaluation of the interpretability of detected relationships

To assess the extent to which the extracted relationships could be explained by existing knowledge, we randomly sampled 100 of the 1,824 detected relationships and examined their interpretability. As a result, 65% (65/100) could be explained on the basis of existing biological knowledge (Table S4). Stratified by correlation strength, 100% (4/4) of very strong correlations, 84% (16/19) of strong correlations, and even 58% (45/77) of moderate correlations were explainable (Figure 2C). These results indicate that stronger correlations tend to be easier to map onto known mechanisms, while even moderate correlations contain substantial mechanistic information.

By contrast, 35% of the relationships were difficult to explain with current knowledge. However, this does not mean that they are meaningless correlations. Indeed, the shuffle control produced almost no incidental high correlations. Unexplained relationships may include those arising from as-yet-uncharacterized mechanisms, those mediated by confounding or undefined intermediate factors, or those involving later developmental stages for which knowledge remains limited.

Although the developmental-stage difference did not significantly differ between the explainable and unexplainable groups (p = 0.08; Figure 2D), a significant difference was observed in the distribution of the later developmental stage within each character pair (p = 0.003; Figure 2E). This suggests that limited knowledge of later developmental processes is one factor underlying the difficulty of interpretation.

### Construction of a phenotypic expression network

To understand the extracted relationships as a whole, we examined the connectivity among phenotypic characters induced by the detected correlations. Of the 421 defined phenotypic characters, 92% (388/421) were connected to at least one other character by a correlation. For these 388 characters, we assigned 46 developmental stages as the temporal axis and cell-wise categories as the spatial axis, and visualized the result as a network in which the edges represent relationships between phenotypic characters (Figure 3A). This network provides a coarse-grained map of which characters tend to co-vary with characters at particular times and in particular cells, and can therefore be viewed as a spatiotemporal phenotypic expression network in early embryogenesis. Because 89% (346/388) of the connected characters formed a single giant connected component, many phenotypes in early embryogenesis appear to be densely interconnected.

**Figure 3.**
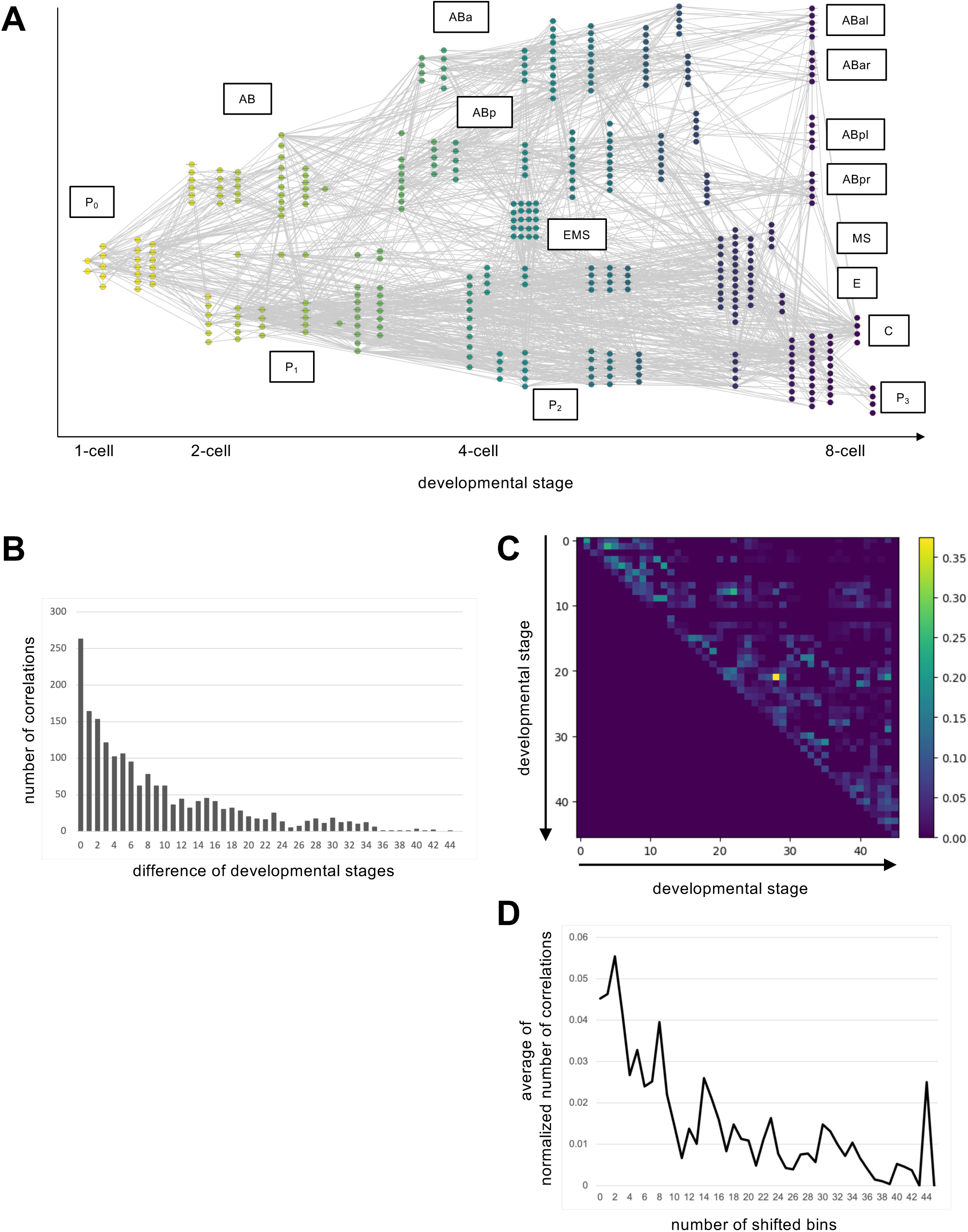
Construction and analysis of the phenotypic expression network. (A) A network of phenotypic characters organized by developmental time and cell identity, where nodes represent phenotypic characters and edges represent correlations. Time was classified into 46 developmental stages, and space was classified by cell. (B) Histogram of developmental-stage differences for character pairs linked by correlations. (C) Heat map showing connectivity across the 46 developmental stages. Values were normalized by dividing the number of detected correlations by the number of characters defined in the corresponding stages. (D) Mean normalized connectivity as a function of stage difference from the diagonal.

When we examined the 33 characters for which no correlations were detected, 39% (13/33) were defined as differences from the wild-type mean and were concentrated near zero. For characters with an extremely narrow value range, statistical detection of relationships with other characters becomes difficult. In addition, 30% (10/33) were characters related to spindle width or length, which are likely to be sensitive to region-detection errors that depend on image quality. Thus, the absence of a detected correlation does not immediately imply biological independence; it also reflects reduced statistical power caused by restricted value ranges or measurement noise.

### Analysis of the phenotypic expression network

To examine the temporal range over which early phenotypes may influence later phenotypes, we analyzed the differences in developmental stage between character pairs linked by a correlation. First, for all correlations, we created a histogram of the differences between the 46 developmental stages. Relationships within the same stage were most abundant, and the number of relationships decreased as the stage difference increased (Figure 3B). This suggests that characters measured at nearby developmental times are more densely related.

Next, to correct for differences in the number of characters defined at each developmental stage, we normalized the number of detected relationships between two stages by the number of characters defined at those stages. This yielded a heat map showing connectivity across the 46 developmental stages (Figure 3C). Values were high near the diagonal and decreased with increasing temporal distance, and this trend was also reproduced when all correlation coefficients were used rather than only thresholded relationships (Figure 3D, Figure S1). Together, these results suggest that phenotypic characters can influence the expression of later characters, but that the temporal range of this influence is relatively limited, with development proceeding as a chain of relationships between temporally adjacent states.

### Extraction of candidate genes based on relationship-disrupting outliers

Next, to extract genes involved in the mechanisms supporting the relationships observed in the wild type, we developed an outlier-analysis framework using quantitative data under perturbation conditions. If a gene involved in a process supporting a particular correlation is perturbed, that process is expected to fail and the relationship between phenotypic characters maintained in the wild type is expected to collapse. In such cases, the perturbed sample is observed as a relationship-disrupting outlier (Figure 1B). In this framework, for each character pair detected in the wild type, we determined whether samples under perturbation conditions fell outside the prediction interval and extracted genes that support the relationship as mechanistic candidate genes.

This determination used prediction intervals estimated from wild-type data. A prediction interval specifies the range within which a future observation is expected to fall with a given probability, conditional on the already observed character value. If a perturbed sample falls outside this prediction interval, the perturbation is interpreted as having induced a state in which the dependence between the two characters maintained in the wild type can no longer be preserved. Thus, the framework prioritizes genes that selectively disrupt specific relationships as mechanistic candidates.

### Detection of candidate genes in real data

To examine whether the framework can detect relationship-disrupting outliers in real data, we applied it to quantitative data on nuclear division dynamics during early embryogenesis in RNAi embryos of *C. elegans*. First, we extracted 421 phenotypic characters from 1,142 RNAi embryo datasets targeting 263 embryogenesis-essential genes [14] and obtained phenotypic expression profiles for the RNAi embryos. We then applied the framework to the relationships detected from wild-type data and evaluated for which relationships each RNAi embryo became an outlier. For character pairs spanning different developmental stages, the earlier character was used as the explanatory variable and the later character as the response variable; for character pairs within the same stage, both directions were evaluated.

Of the 1,824 detected correlations, 263 were identified within the same developmental stage; therefore, including bidirectional evaluation, outlier detection was performed for a total of 2,087 character pairs. As a result, 129,100 outlier instances were detected (Table S5). When cases in which the same relationship was an outlier in multiple embryos targeting the same gene were collapsed, these were reduced to 98,922 gene-relationship correspondences. Furthermore, 99.6% (262/263) of the analyzed genes were detected as outliers for at least one relationship (Figure 4A; Table S6). Genes ranked high by the number of detected outlier relationships were enriched for functions related to astral microtubules, spindle-axis formation, and mitotic spindle assembly (Table S7).

**Figure 4.**
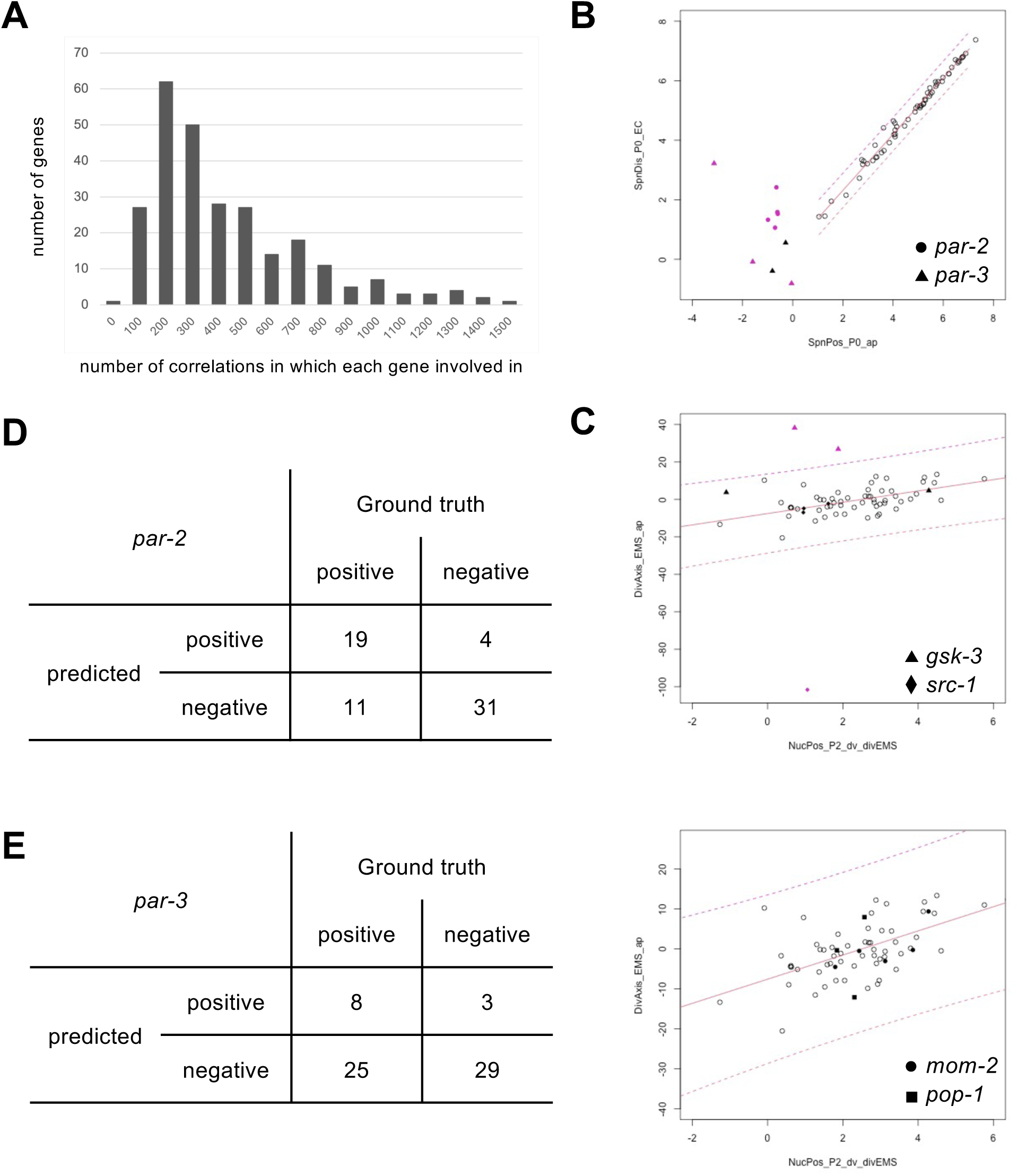
Candidate-gene extraction based on relationship-disrupting outliers. (A) Number of correlations for which each gene was detected as an outlier. (B-C) Validation using representative mechanistic examples. (B) *par-2* and *par-3* RNAi embryos were detected as outliers for the relationship corresponding to the establishment and maintenance of cell polarity. (C) *gsk-3* and *src-1* RNAi embryos were detected as outliers for the relationship corresponding to EMS spindle orientation, whereas *mom-2* and *pop-1* RNAi embryos were not. (D) Performance evaluation for *par-2*. (E) Performance evaluation for *par-3*.

### Validation of candidate-gene extraction

To show that the framework preferentially detects genes involved in known mechanisms, we examined two representative mechanistic examples. As the first example, we focused on the mechanism that positions the spindle in the P_0_ cell. Formation of the P_0_ spindle along the anterior-posterior axis and its displacement toward the posterior pole are achieved through the establishment and maintenance of cell polarity dependent on PAR-2 and PAR-3 [19]. Indeed, when RNAi embryos targeting *par-2* and *par-3* were mapped onto the relevant relationship, they were detected as relationship-disrupting outliers for the relationship corresponding to the P_0_ spindle-positioning mechanism (Figure 4B). These results show that the framework can appropriately recover genes involved in the known polarity-establishment mechanism.

As the second example, we focused on the mechanism supporting EMS spindle orientation. This spindle orientation is known to be regulated by the Wnt pathway involving MOM-1, MOM-2, MOM-5, KIN-19, and GSK-3, together with the parallel SRC-1/MES-1 pathway [23,24,25]. By contrast, APR-1, WRM-1, and POP-1 are involved in endoderm induction but are not thought to be directly involved in EMS spindle orientation itself. We therefore evaluated RNAi embryos targeting *mom-2*, *gsk-3*, *src-1*, and *pop-1*. RNAi embryos targeting *gsk-3* and *src-1* were detected as outliers for the relevant relationship, consistent with the known involvement of the Wnt and SRC-1/MES-1 pathways. By contrast, *pop-1* RNAi embryos were not detected as outliers, consistent with the established view that *pop-1* is involved in endoderm induction rather than spindle orientation. *mom-2* was also not detected as an outlier, but this is consistent with the report of Schlesinger et al. [24], who showed that *mom-2/wg* is not essential for EMS spindle orientation because a parallel signal can compensate (Figure 4C). Together, these results show that the framework can extract, as relationship-disrupting outliers, genes involved in the mechanisms supporting relationships between phenotypic characters.

### Evaluation of precision and recall for candidate-gene extraction

To evaluate the performance of the candidate-gene extraction framework, we examined whether *par-2* and *par-3*, whose functions are well characterized [26], were detected as outliers for relationships in which they were expected to be involved. Specifically, among the 100 relationships randomly sampled above, we focused on the 65 relationships judged to have an underlying mechanism, classified from existing knowledge whether each gene was expected to function in that mechanism, and used these classifications as gold-standard labels to calculate precision and recall (Figure 4D, 4E; Table S9).

For *par-2*, 23 of the 65 relationships were detected as outliers. Of these 23 relationships, 19 were also expected to involve *par-2* according to the gold-standard labels based on existing knowledge, yielding a precision of 83% (19/23). Because *par-2* involvement was expected for 30 relationships in the gold-standard labels, the recall was 63% (19/30). For *par-2*, both precision and recall were relatively high.

A similar evaluation for *par-3* yielded a high precision of 73% (8/11) but a low recall of 24% (8/33). Thus, while the framework is effective for narrowing down candidate genes, exhaustive recovery of all truly involved relationships still depends on experimental factors such as RNAi penetrance, phenotypic pleiotropy, and the sensitivity of the evaluated characters. Even so, the high precision indicates that the framework is useful for prioritizing mechanistic candidate genes.

## Discussion

The main focus of this study is not a comprehensive description of nematode embryogenesis itself, but the presentation of an analytical framework applicable to quantitative data of morphological dynamics in general. Nuclear division dynamics during early embryogenesis in *C. elegans* is positioned here as a proof-of-concept system for validating the framework. Recent advances, including deep-learning methods that reconstruct whole-embryo cell lineages from sparse annotations and methods that track cell lineages from three-dimensional time-series images, are rapidly lowering the barrier to preparing quantitative data of morphological dynamics for control and perturbation groups [27,28]. The framework should therefore be extensible to diverse systems, including developmental processes, cultured cells, and organoids.

This framework does not equate correlation with causation, nor is it a formal causal-inference algorithm. This distinction should be stated explicitly. However, that is not the limit of the present claim. Relationships between phenotypic characters that are reproducibly observed in quantitative data of morphological dynamics, especially when they are selectively disrupted under perturbation conditions, contain mechanistic information beyond simple co-occurrence statistics. Multiple generative mechanisms can underlie such relationships, including direct causation, indirect causation, common upstream factors, geometric constraints, shared variables arising from character definitions, and latent confounding; nevertheless, in every case they represent biologically grounded relationships that warrant experimental testing. Indeed, 65% of the randomly sampled relationships could be explained by existing knowledge, and the remaining 35% are unlikely to be mere products of chance in light of the shuffle control. Thus, although the relationships extracted by the framework do not constitute final proof of causality, they are meaningful as high-value mechanistic candidates that merit experimental validation.

In the current implementation, linear correlation was used as the dependency metric and prediction intervals from linear regression were used for outlier detection. This choice allowed large-scale exploration of hundreds of characters and thousands of relationships while retaining interpretability. However, the current implementation can be extended to capture nonlinear relationships, accommodate changes in variance across the data range, and use models that are robust to outliers. In the future, combining MIC [29], distance correlation [30], Lasso regression [31], and LiNGAM [32] may allow the framework to handle more diverse relationship structures and candidate causal directions.

The phenotypic expression network constructed from correlations among phenotypic characters provided a useful coarse-grained model of spatiotemporal relationships in early embryogenesis. That 346 characters formed a single giant connected component indicates that early embryogenesis proceeds as an integrated process composed of many interlinked characters. In addition, because normalized connectivity based on correlations declined as temporal distance increased, development is more likely to proceed through chains of relationships between nearby time points than through long-range direct dependencies. Such network representations are useful for grasping the global structure of development, which is difficult to infer from scatter plots of individual character pairs alone.

Interpretation of candidate genes must be made with RNAi penetrance, sample size, and phenotypic pleiotropy in mind. Even when a gene truly participates in a mechanism, outliers may be confined to only a subset of samples if perturbation strength is heterogeneous. Conversely, strong upstream perturbations or widespread morphological abnormalities can generate secondary outliers across many relationships. Indeed, genes with large numbers of outliers were enriched for functions in spindle formation and cytoskeletal processes, suggesting that the framework can read out the scope of upstream mechanisms from the distribution of relationship disruption itself. Outlier detection should therefore be interpreted not as a simple binary judgment but together with detection frequency and spatiotemporal distribution for each relationship.

### Conclusion

We developed a computational framework that extracts reproducibly observed relationships between phenotypic characters in wild-type groups from quantitative data of morphological dynamics and evaluates how those relationships are disrupted in perturbed groups. In a proof of concept using nuclear division dynamics during early embryogenesis in *C. elegans*, we showed that the framework can recover relationships between phenotypic characters consistent with known mechanisms and prioritize candidate genes. By reframing correlation as relationship data that carry mechanistic information, this framework provides a basis for systematically generating testable mechanistic hypotheses.

## Materials and Methods

### Quantitative data on nuclear division dynamics

For the proof of concept and performance evaluation in this study, we primarily used a collection of quantitative data on nuclear division dynamics during early embryogenesis in *C. elegans* [14]. This collection consists of quantitative data from 59 wild-type embryos and 1,142 RNAi embryos targeting 263 genes. Each dataset was obtained from time-lapse images of early embryogenesis acquired by differential interference contrast microscopy for 2 h at 20-s intervals and 66 focal planes with a z-step of 0.5016 μm, and measured using a system for nuclear-region detection and tracking [10].

In addition, for several genes used in the validation examples, we performed new RNAi experiments and collected images and quantitative data using the same procedure as in the previous report [14]. In this study, including these additional data, we extracted relationships between phenotypic characters in the wild type and detected relationship-disrupting outliers under perturbation conditions, that is, samples in which a relationship between phenotypic characters maintained in the wild type collapsed under perturbation.

### Detection of correlations between phenotypic characters

To extract candidate relationships between phenotypic characters in wild-type data, we used the Pearson correlation coefficient between two phenotypic characters:

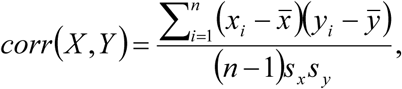

Here, x̅ and y̅ denote the means of phenotypic characters *X* and *Y*, respectively; s_x_ and s_y_ denote the standard deviations; and *n* is the sample size. In this study, character pairs with |*r*| exceeding a predefined threshold were extracted as candidate relationships. Although a highly correlated character pair does not by itself determine a unique causal direction, such a pattern can arise from multiple mechanistic possibilities, including a direct effect, a longer causal chain, a common upstream factor, mechanical or geometric constraints, or unobserved confounding. We therefore treat this multiplicity not as a weakness but as a source of information, and regard highly correlated character pairs as strong candidate relationships that reflect a shared mechanism.

### Detection of relationship-disrupting outliers

To prioritize genes involved in the mechanisms supporting candidate relationships, we constructed prediction intervals for the distribution of wild-type phenotypic characters and evaluated deviations of perturbed samples [33]. The prediction interval was calculated as follows:

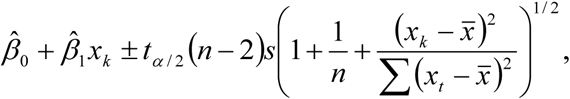

Here, *β*_!_ and *β*_“_ denote the estimated regression coefficients obtained from the wild-type data, *n* is the number of wild-type samples, and *s* is the residual standard error. For character pairs belonging to different developmental stages, the earlier character was used as the explanatory variable and the later character as the response variable. For character pairs within the same developmental stage, both directions were evaluated because no directionality was assumed a priori. The significance level α was set at 0.001.

## Data availability

All relevant datasets are available within the paper and on SSBD:repository (ssbd-repos-000519; https://doi.org/10.24631/ssbd.repos.2026.06.519).

## Acknowledgments

Worm strains were provided by the Caenorhabditis Genetics Center, funded by the NIH National Center for Research Resources. We thank all members of the Onami laboratory for helpful discussions and technical support.

## Author contributions

K.K.: Formal analysis, Methodology, Software, Visualization, Writing – Original Draft; H.O.: Investigation; S.O.: Conceptualization, Supervision, Writing – Review & Editing.

