## Supplementary material for "A Computational Framework for Extracting Mechanistic Hypotheses from Quantitative Data of Morphological Dynamics": Figure S1

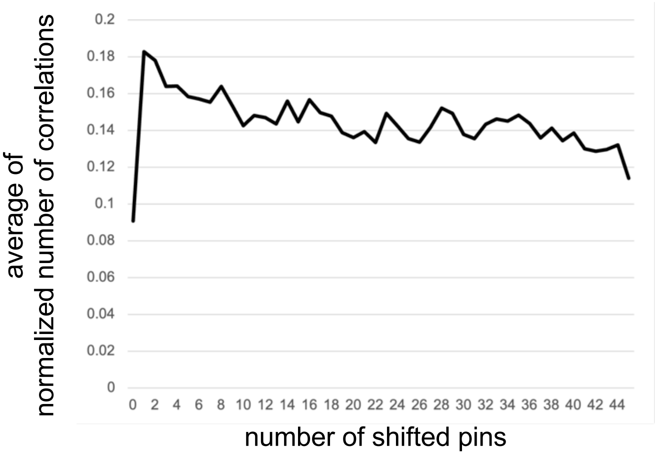


**Figure S1.** Same analysis as in Figure 3D, but using all correlation coefficients instead of the number of detected relationships. For each pair of developmental stages, all correlation coefficients were summed, normalized by the numbers of phenotypic characters assigned to the two stages, and further normalized by the number of stage pairs at each developmental-stage distance. The x- and y-axes are identical to those in Figure 3D.
